# Ultra-deep whole genome bisulfite sequencing reveals a single methylation hotspot in human brain mitochondrial DNA

**DOI:** 10.1101/2021.03.30.437685

**Authors:** Romain Guitton, Christian Dölle, Guido Alves, Ole-Bjørn Tysnes, Gonzalo S. Nido, Charalampos Tzoulis

**Author notes:** To whom correspondence should be addressed.: **Prof. Charalampos Tzoulis**, Neuro-SysMed Center of Excellence for Clinical Research in Neurological Diseases, Department of Neurology, Haukeland University Hospital, Department of Clinical Medicine, University of Bergen, 5021 Bergen, Norway., E-mail-2, **Dr Gonzalo S. Nido**, Neuro-SysMed Center of Excellence for Clinical Research in Neurological Diseases, Department of Neurology, Haukeland University Hospital, Department of Clinical Medicine, University of Bergen, 5021 Bergen, Norway.

## Abstract

While DNA methylation is established as a major regulator of gene expression in the nucleus, the existence of mitochondrial DNA (mtDNA) methylation remains controversial. Here, we characterised the mtDNA methylation landscape in the prefrontal cortex of neurological healthy individuals (n=26) and patients with Parkinson’s disease (n=27), using a combination of whole genome bisulfite sequencing (WGBS) and bisulfite-independent methods. Accurate mtDNA mapping from WGBS data required alignment to an mtDNA reference only, to avoid misalignment to nuclear mitochondrial pseudogenes. Once correctly aligned, WGBS data provided ultra-deep mtDNA coverage (16,723±7,711), and revealed overall very low levels of cytosine methylation. The highest methylation levels (5.49±0.97%) were found on CpG position m.545, located in the heavy-strand promoter 1 region. The m.545 methylation was validated using a combination of methylation-sensitive DNA digestion and quantitative PCR analysis. We detected no association between mtDNA methylation profile and Parkinson’s disease. Interestingly, m.545 methylation correlated with the levels of mtDNA transcripts, suggesting a putative role in regulating mtDNA gene expression. In addition, we propose a robust framework for methylation analysis of mtDNA from WGBS data, which is less prone to false-positive findings due to misalignment of nuclear mitochondrial pseudogene sequences.

**Graphical abstract of the analyses and main findings:** Fresh-frozen brain tissue was obtained from the prefrontal cortex (Brodmann area 9) of 53 individuals, comprising 27 patients with idiopathic PD and 26 healthy controls. Tissue from the same samples was used in three different downstream analyses. WGBS was conducted on all 53 samples and the data were analysed using three different alignment strategies. Alignment against an mtDNA reference only was clearly superior as it gave the highest and most even depth of coverage. WGBS analysis revealed that mtDNA harbours very low levels of cytosine methylation, with the exception of the CpG position m.545 within the HSP1 region (lower right inset). The m.545 methylation was confirmed by bisulfite- and sequencing-independent methods, employing methylation-specific MspJI DNA digestion, followed by quantification with qPCR or fluorescent PCR and capillary electrophoresis. Finally, mtDNA transcript levels were determined by RT-qPCR and correlated to m.545 methylation levels, showing a positive association.

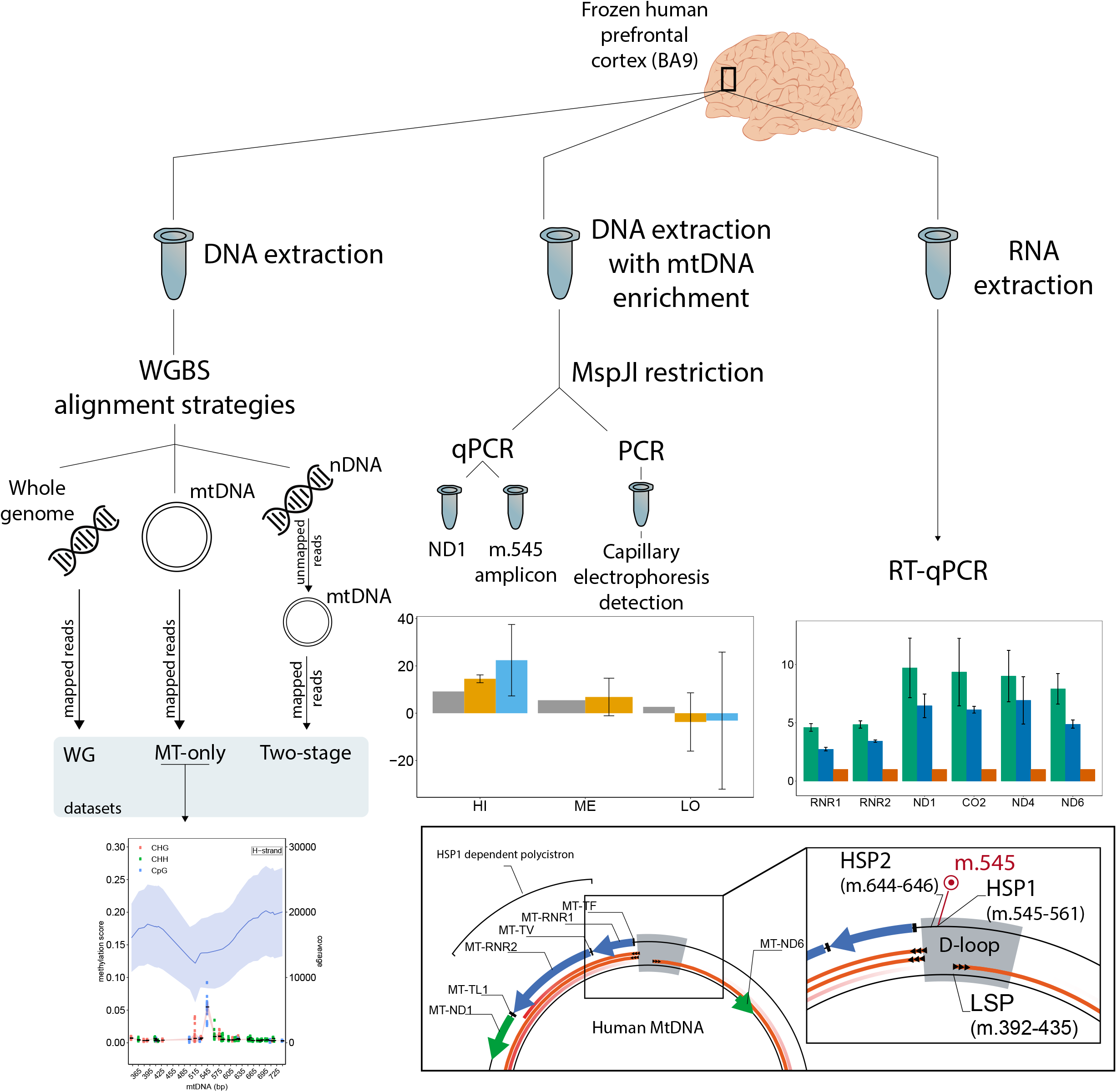

## INTRODUCTION

The human mitochondrial DNA (mtDNA) is a double-stranded, 16,569 base pairs (bp) long, circular genome, which is present in multiple copies per cell. mtDNA encodes a total of 37 genes: 13 peptide subunits of the respiratory chain complexes I, III, IV and V, as well as 22 transfer RNAs (tRNAs) and two ribosomal RNAs (rRNAs), which are required for the translation of the peptide genes. All remaining mitochondrial proteins are encoded in the nuclear genome. The two strands of mtDNA have been traditionally termed heavy (H-) and light (L-) strand, due to different C/G content. The majority of genes (28 genes) are encoded on the L-strand (i.e. transcribed from the H-strand), while 9 are encoded on the H-strand (1). mtDNA has a single ~1,200 bp long regulatory region (m.16,024-576), which contains the displacement loop (D-loop), the origin of replication of the H-strand, and the promoters and origin of transcription for both strands (2, 3). The L-strand promoter (LSP) controls transcription of MT-ND6 and eight tRNA genes. Two promoters have been identified for the H-strand: HSP1 and HSP2. Current evidence suggests that HSP1 (m.545-567) regulates the expression of two rRNA and two tRNA genes, while HSP2 (m.644-646) regulates the transcription of the remaining 12 protein coding and 14 tRNA genes of the H-strand. However, the exact number and function of the mtDNA H-strand promoters remains debated (4, 5).

mtDNA encoded genes are essential for the assembly and function of the respiratory chain and their mutations cause a broad spectrum of monogenic diseases, affecting approximately 1:5,000 individuals (6). Moreover, both inherited variation and somatic mtDNA changes have been associated with susceptibility to Parkinson’s disease (PD) (7, 8) and other neurodegenerative disorders (9, 10), as well as with the process of aging (11). While mtDNA genetics has been extensively studied in health and disease, the role of epigenetic regulation of the mitochondrial genome remains largely unexplored. Several studies have reported evidence of mtDNA methylation, with varying levels and position (12–18), and some have suggested that mtDNA methylation may be altered in neurodegenerative diseases, such as PD and Alzheimer’s disease (AD) (19, 20). Others, however, reported no evidence that mtDNA methylation occurs at all (21–28). These conflicting findings may be partly confounded by technical artifacts and limitations applying specifically to the study of mtDNA. First, it has been suggested that, unless linearised, the coiled structure of mtDNA may prevent bisulfite conversion, resulting in false positive methylation results (29). Second, mapping to mtDNA is prone to misalignment artifacts caused by highly similar *nuclear mitochondrial DNA sequences* (NUMTs) (30, 31). Since cytosines are often methylated in nuclear DNA, NUMT-misalignment to mtDNA may generate false positive methylation findings. Thus, the questions of whether mtDNA methylation occurs and whether it plays a role in aging and disease, remain unresolved.

Here, we assess mtDNA methylation at high resolution, using ultra-deep whole genome bisulfite sequencing (WGBS) data from prefrontal cortex tissue of 27 individuals with PD and 26 healthy controls. We show that mtDNA harbours very low levels of cytosine methylation, with the exception of the CpG position m.545 within the HSP1 region. We confirm this finding, using a combination of methylation-sensitive DNA digestion and quantitative PCR. Furthermore, our analyses suggest that the levels of the m545 methylation increase slightly with age and correlate to mtDNA gene expression levels, but show no association to PD. Finally, our findings help establish an accurate framework for studying mtDNA methylation from WGBS data.

## MATERIAL AND METHODS

### Subject cohorts

We studied fresh-frozen prefrontal cortex (Brodmann area 9) tissue from a total of 53 individuals from two independent cohorts. One cohort (n = 32) comprised individuals with idiopathic PD (n = 17) from the Norwegian Park-West study (PW), a prospective population-based cohort (32) and neurologically healthy controls (Ctrl, n = 15) from our brain bank. The controls comprised a group that was demographically matched with the PD patients (n = 11) and a younger group (n = 4) of children aged from 0 to 0.38 years old. Whole-exome sequencing had been performed on all patients (33) and known/predicted pathogenic mutations in genes implicated in Mendelian PD and other monogenic neurological disorders had been excluded. None of the study participants had clinical signs of mitochondrial disease or use of medication known to influence mitochondrial function. The second cohort comprised individuals with idiopathic PD (n = 10) and demographically matched neurologically healthy controls (n = 11) from the Netherlands Brain Bank (NBB). Individuals with PD fulfilled the National Institute of Neurological Disorders and Stroke (34) and the UK Parkinson’s disease Society Brain Bank (35) diagnostic criteria for the disease at their final visit. A detailed description of the study participants is provided in Supplementary Table S1.

Whole genome bisulfite sequencing (WGBS), quality control and comparison of alignment strategies

Extraction of DNA, sonication and bisulfite sequencing (aiming at 30X coverage) was conducted by Hudson-Alpha Discovery (Huntsville, AL). Sonication allows DNA linearisation of circular DNA and facilitates bisulfite action on cytosines as reported (29). A methylation-free DNA fragment (Lambda phage spike-in) was included in each sample to assess the bisulfite conversion efficiency.

Quality control of the raw sequence files was carried out using fastQC (36) before read trimming. Final trimming settings were determined following M-bias estimation using Bismark 0.17.0 (37) and performed using Trimmomatic version 0.33 (38): “ILLUMINACLIP:TruSeq3-PE-2.fa:2:30:10 CROP:145 HEADCROP:10 LEADING:3 TRAILING:3 SLIDINGWINDOW:4:15 MINLEN:30”. The range of read lengths after trimming was 30-135bp. Quality control and M-bias of the resulting trimmed reads was then reassessed.

Sequence alignment was performed by Bismark 0.17.0 employing three alternative strategies. In the first strategy, reads were aligned to the whole GRCh38 reference genome (39), including the rCRS mitochondrial reference (40). In the second strategy (“MT-only”), reads were aligned against the rCRS (40) sequence only. Finally, in the third strategy (“two-stage mapping”) reads were mapped against the nuclear genome first (excluding the mitochondrial chromosome), and unmapped reads were subsequently mapped against the rCRS reference. These alternative approaches were performed to evaluate the effects of sequence similarities between mtDNA and nuclear DNA, specifically “Nuclear mitochondrial DNA” (or NUMTs) (30, 31). Alignment files resulting from each of the three strategies were deduplicated using Bismark deduplicate and per-base methylation call performed using Bismark methylation extractor, both part of Bismark 0.17.0 software.

The mean coverage amongst samples per position was then calculated and the mean coverage per 100-bp region averaged over all positions in this region. In order to explain the differences between the alignment strategies, we extracted the positions defined by a coverage lower than 1,000 in the whole-genome alignment. We assessed coverage discrepancies across the three alternative mapping strategies by tracing the positions of individual mapped reads using their unique ids. Methylation per read, mapping quality and mapping position were extracted with Bismark and compared to the first strategy, which was used as a reference. Depth of coverage on extra-mitochondrial regions was assessed using Samtools 1.9 (41). An additional probabilistic approach was used to study the general discrepancies in mapping between the alignment strategies. We randomly downsampled to 300,000 reads per sample mapped with the MT-only strategy and were traced in WG-alignment. Subsequently, cytosines (both methylated and unmethylated) were numbered and methylation ratio per read was calculated.

### Methylation analysis

The methylation score was calculated for each cytosine position in the reference genome as the ratio of counts of C to the sum of counts of C and T. To ensure a high confidence in our analysis, we discarded positions with 0 depth in more than 95% of the samples in our cohort. This rule applies also when a minimum depth threshold was used. The minimum depth threshold was used as a complementary filter and set to 1,000 reads, as we sought to call methylation scores down to 1% with at least 10 supporting reads on the minor allele. Finally, to minimise the chance of false positive methylation calls, and given the average read length, we excluded from the analysis the first and last 125 positions of the mitochondrial genome. Methylation score and coverage for a given position are reported as the mean over all samples for which the position is covered by at least one read, with the standard deviation reported unless specified otherwise. Rolling means for coverage and methylation were calculated with an order 3 (m=3), i.e. taking the mean of the considered cytosine and the two preceding defined cytosines.

We analysed cytosine methylation relative to its context with the convention: CpG, CHH or CHG, where H stands for a non-guanine nucleotide.

We tested for batch effects using Spearman correlation test for continuous data and Kruskal-Wallis or Mann-Whitney-Wilcoxon tests for discrete data using R (42); the significance level was set to 0.05.

### mtDNA extraction and enrichment

Fresh-frozen brain tissue samples (Brodmann area 9) adjacent to those used for WGBS were analysed. To enrich for mitochondria, the tissue was lysed in 300 μl of a mannitol-sucrose buffer (225 mM Mannitol, 75 mM sucrose, 10 mM Tris-HCl pH 7.6, 1 mM EDTA, 0.1 % fatty-acid free BSA) using a Dounce homogenizer. The pestle was rinsed with 700 μl of the same solution and both solutions combined. The homogenate (1 ml total) was centrifuged at 800× g and 4 °C for five minutes. The supernatant was kept on ice in a new tube while the pellet underwent the previous steps for a second time and both supernatants were combined. Samples were centrifuged at 5,000× g and 4 °C for ten minutes. The residual pellet was resuspended in 180 μl of buffer ATL (QIAGEN DNeasy blood & tissue kit) and digested by proteinase K overnight (1 μl per 1 mg tissue) at 56 °C. 200 μl of buffer AL (QIAGEN) was added followed by ten minutes of incubation at 70 °C to deactivate proteinase K. The tube was centrifuged at 13,000× g and 4 °C for five minutes, the supernatant removed; the pellet washed in 1 ml of the mannitol-sucrose buffer and centrifuged again. The pellet was resuspended in 500 μl of water and the contained DNA precipitated by adding 30 μl 5M NaCl, 3 μl glycogen (Thermo Scientific R0561) and 530 μl isopropanol, and gently inverting six times as described (43), followed by incubation at −20 °C overnight. Centrifugation at 14,000 rpm and 4 °C for 60 minutes yielded a pellet that was washed with 1 mL 70% EtOH and resuspended in 30 μl water. We measured the DNA concentration and stored the sample at −20 °C until further use.

### MspJI restriction digestion and hybridisation of synthetic dsDNA oligonucleotides

DNA fragments were incubated with MspJI (NEB R0661) for four hours at 37 °C following the manufacturer’s recommendation. MspJI recognises the sequence ^m^CNNR and exclusively digests the DNA if the cytosine is methylated (44, 45).

In order to test the suitability of our approach, which relied on the ability of MspJI to distinguish between methylated and non-methylated cytosine at position m.545 on the heavy-strand of mtDNA, we designed single-strand DNA oligonucleotides (Supplementary Table S2) and hybridised them in several combinations to mimic the area of interest (m505-603) in non-methylated, hemi-methylated and fully methylated states with regard to the MspJI restriction site. Hybridisation was achieved by incubating equimolar amounts of two single-stranded, complementary oligonucleotides for five minutes at 95 °C followed by cooling stepwise to 10 °C. Samples were stored at −20 °C until use. Methylation sensitive PCR amplification Quantitative PCR (qPCR) and fragment analyses measured the differential amplification of the same sample incubated with and without MspJI. MspJI cleaved the mtDNA sequence when m.545 was methylated, which reduced the PCR amplification of this region accordingly. The primers and probes for qPCR are shown in Supplementary Table S2 and the following conditions were used: 2 μl (~15 ng) digested or non-digested mtDNA sample was used as template per reaction and all samples were run in triplicates. Amplification was performed on a StepOne Plus instrument (Applied Biosystems) using TaqMan Fast Advanced Master Mix (Thermofisher). Thermal cycling consisted of one cycle at 95□°C for 20□seconds and 40 cycles at 95□°C for three seconds and 60□°C for 30□seconds. Using the 2^-Δ/ΔCT^ Livak method (46), we calculated the ratio of the amplicon to the reference (MT-ND1) and compared the amplification from digested and undigested template following the formula:

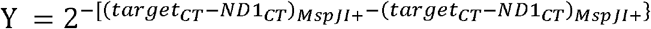

is amplification ratio value was subtracted from 1 in order to retrieve the m.545 methylation score in percent.

For fragment analysis, the primer sequences were identical to the qPCR approach, but primers were fluorescently labelled. PCR amplification was carried out as for qPCR analysis, albeit for 30 cycles. Samples were analysed using capillary electrophoresis on a ThermoFisher Veriti sequencer instrument. We filtered for fragment size (61 to 65 bp as the expected size was 64 bp) and peak height (above 500 units intensity) to avoid background contamination. The peak area was measured and the ratio between samples incubated with and without MspJI calculated as the relative abundance of unmethylated DNA. This ratio subtracted from 1 provided the m.545 methylation score.

### RNA extraction

RNA was extracted from fresh-frozen brain tissue (Brodmann area 9) using the RNeasy lipid tissue mini kit (QIAGEN); 20 mg of tissue was homogenised in 1 mL QIAzol lysis reagent using a 1 mL syringe successively with 23G and 25G needles. The homogenate was incubated for five minutes at room temperature and then 200 μl chloroform was added. After a vigorous shake, the samples were incubated for three minutes at room temperature and centrifuged at 12,000× g for 15 min at 4 °C. The aqueous phase was transferred into a new tube and the same volume of 70 % EtOH added. The sample was vortexed and the mixture applied to an RNeasy mini spin column and centrifuged at 8,000× g for 15 seconds. Subsequently, 700 μl buffer RW1 was added and the column centrifuged at 8,000× g for 15 seconds. The RPE buffer was added in two steps, separated by a centrifugation at 8,000× g for 15 seconds, and then finally centrifuged at 8,000× g for two minutes. RNA was eluted in 30 μl RNase-free water by centrifugation at 8,000× g for one minute. For concentration determination, 1 μl was taken out and diluted to be measured in a Nanodrop, and the sample stored at −80 °C until further use.

Complementary DNA (cDNA) synthesis was carried out with the SuperScript IV VILO Kit (ThermoFisher), using 1 μg total RNA as template according to the manufacturer’s protocol.

### qPCR and estimation of mtDNA copy number per cell

The following Taqman assays (ThermoFisher) were used for cDNA quantification of a selection of genes from the catalogue (Supplementary Table S2). For *MT-ND4*, we used custom-designed primers and probe (Supplementary Table S2).

*GAPDH* was used as reference and relative quantification was performed using the 2^-(ΔCT)^ formula. Thermal cycling was as described above.

Estimation of mtDNA copy number per cell was carried out based on WGBS data adapted from a previous study (47). We calculated the ratio of the average of mtDNA coverage to the average nuclear DNA coverage of chromosome 1. We then multiplied this ratio by two to account for the two copies of nuclear DNA in each cell.

## RESULTS

### Accurate mtDNA sequence alignment from WGBS data requires mapping to the mitochondrial reference genome

It has been shown that mapping to a mtDNA reference from whole-genome data is prone to misalignment due to NUMTs (30, 31). To establish an accurate mapping method for mtDNA, we compared three different alignment approaches: 1) against the whole human genome (WG), 2) against mtDNA only (MT-only), and 3) sequentially against the nuclear genome and mtDNA (two-stage). In this approach, we identified unmapped reads after aligning against the nuclear genome and mapped these against the mtDNA reference. The MT-only alignment produced an even coverage with a mean depth of 16,723±7,711 (range: 4,535–37,696) (Figure 1, Supplementary Table S3). In contrast, the WG-alignment showed lower mean depth (15,192±7,952, range 79–36,105) and uneven mtDNA coverage with a near-zero coverage drop around the region m.4,400-4,700 for the L-strand (Figure 1). The two-stage approach showed even lower mean depth (8,207±8,656, range 0–33,981) and highly uneven coverage with marked decrease in depth, particularly pronounced in the region m.4,000-9,500 on both strands (Figure 1). Estimated methylation was generally very low and uniformly distributed along the entire mtDNA reference in the MT-only approach and in regions of high-coverage in the WG and two-stage alignments. Regions with coverage defects, however, showed a markedly uneven distribution of estimated methylation with prominent deviations from the surrounding sequence, a higher dispersion, and a tendency to overestimate methylation levels (Figure 1).

**Figure 1.**
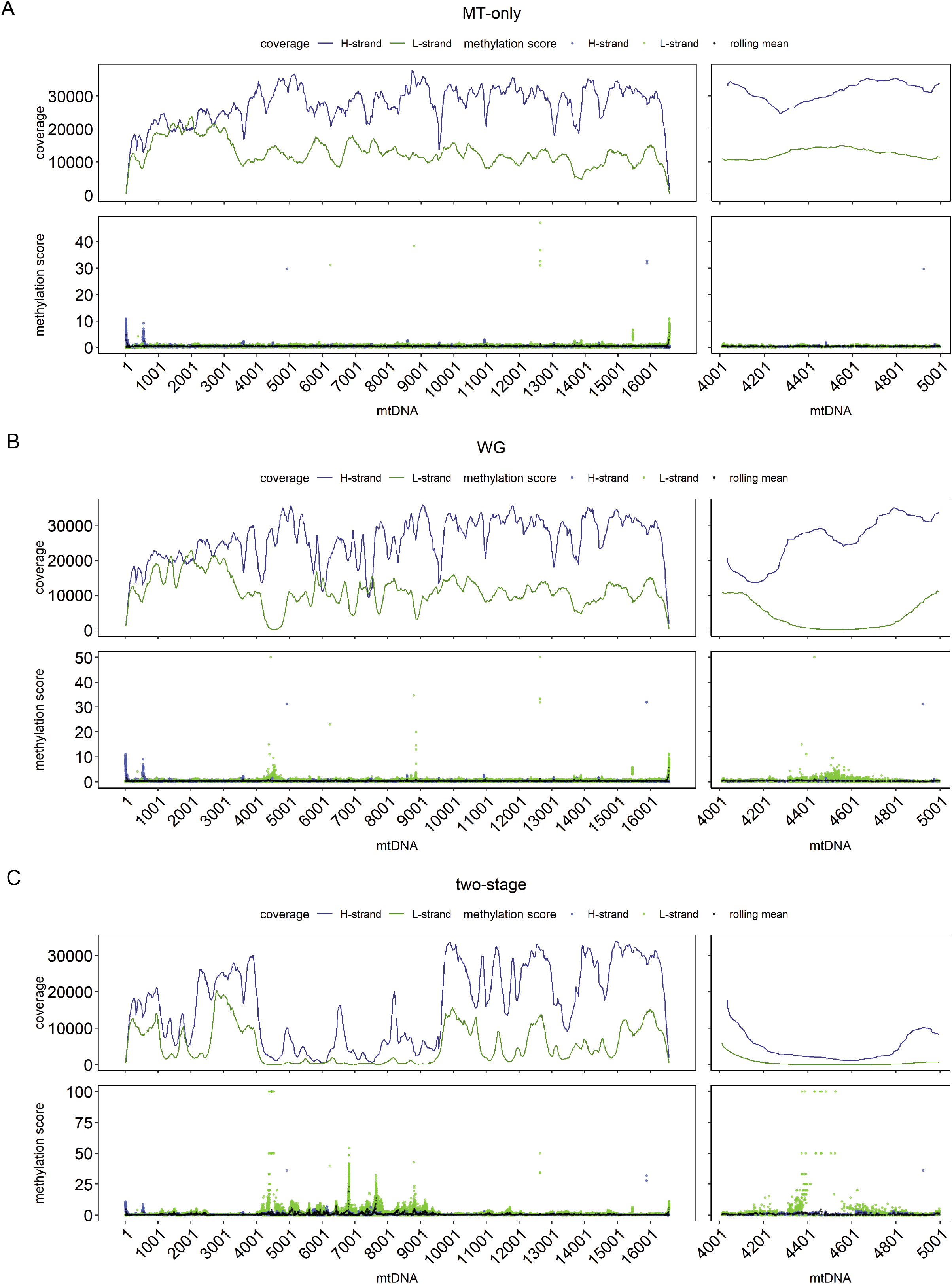
mtDNA depth of coverage and methylation with three different alignment strategies. The plots show depth of coverage and methylation score obtained by the three alignment strategies: A) MT-only, B) WG and C) two-stage. In each alignment approach, the upper panels show the rolling mean depth of coverage (m=3) for the L-strand (green) and H-strand (blue) of the mtDNA. The lower panels show the rolling mean methylation (m=3, black dots) and mean methylation of all samples for each position on the L-strand (green) and H-strand (blue). Data is shown for the whole mtDNA molecule (left) and a magnified region m.4,000-5,000 (right).

The presence of mtDNA regions with low-depth, when the nuclear genome was included in the mapping reference (WG- and two-stage alignments), indicated a possible misalignment on the mtDNA reference due to NUMT sequences. To investigate this, we identified in the WG-alignment all positions supported by less than 1,000 reads and showing lower coverage compared to the MT-only alignment and selected three samples among those with the highest discrepancies. Next, we extracted the reads supporting these positions and traced where they had mapped in the MT-only and WG approach. In the MT-alignment, all reads were mapped to the mtDNA gene *MT-ND2*, encoding the ND2 subunit of respiratory complex I. In the WG-alignment, however, 88.2±0.46 % of these reads (7,120/8,082) had been mapped to the telomeric region of the short arm of chromosome 1 (chr1:629,500-630,000, Figure 2, Supplementary Figure S1) on *MTND2P28* (HGNC:42129), a pseudogene with high sequence similarity to *MT-ND2*. As a result of the misalignment of mtDNA reads, this region of chromosome 1 showed an unusually high depth (2,625±1,089). In comparison the mean sequence depth of chromosome 1 was 35±17. Notably, the mean mapping quality (MAPQ) of the analysed reads was lower when mapping to chromosome 1 (24±9.6) than mapping to mtDNA (37±8.4). However, the MAPQ alone cannot guide the choice of mapping strategy as 69.6±3.7% of the misaligned reads had a MAPQ > 30 (Supplementary Figure S2). Additionally, we randomly chose a subset of 300,000 reads from the whole dataset that had been aligned with the MT-only approach in each of the three samples (900,000 reads total) and investigated their position and characteristics in the WG-alignment. In accordance with what we have shown, 4,266 reads (0.47±0.15%) mapped to the homologous region of the chromosome 1 and showed a low mean methylation of 0.86±0.81%, in line with a mtDNA origin. A total of 276 reads (0.03±0.004%) mapped to chromosomes 2, 3, 5, 6, 11, 13, 14, 17 and X, and presented a mean methylation of 7.4±2.5% (Supplementary Table S4). These observations strongly suggest that reads originating from the mtDNA are erroneously mapped to a homologous region in chromosome 1 when this region is available as a reference at the mapping stage. Read mapping on other chromosomes is notably lower, and their relatively higher methylation levels may indicate a combination of both mitochondrial and nuclear reads. Extending these analyses, we assessed the discrepancy in coverage for all positions between the MT-only and WG alignments. The results showed that WG had a systematically lower coverage with similar depth and methylation score between all cytosine contexts (Supplementary Figure S3A), but a clear difference between strands (Supplementary Figure S3B). The same analysis for the two-stage alignment strategy showed an even larger difference in coverage from the MT-only alignment (Supplementary Figure S3C), further implying that reads originating from mtDNA are prone to erroneously map to nuclear mitochondrial pseudogenes, thereby increasing the dispersion and potentially biasing methylation estimates. Our findings indicate that, in order to avoid this bias and minimise the loss of reads originating from the mtDNA to the nucleus, WGBS data should be aligned to the mtDNA reference only.

**Figure 2.**
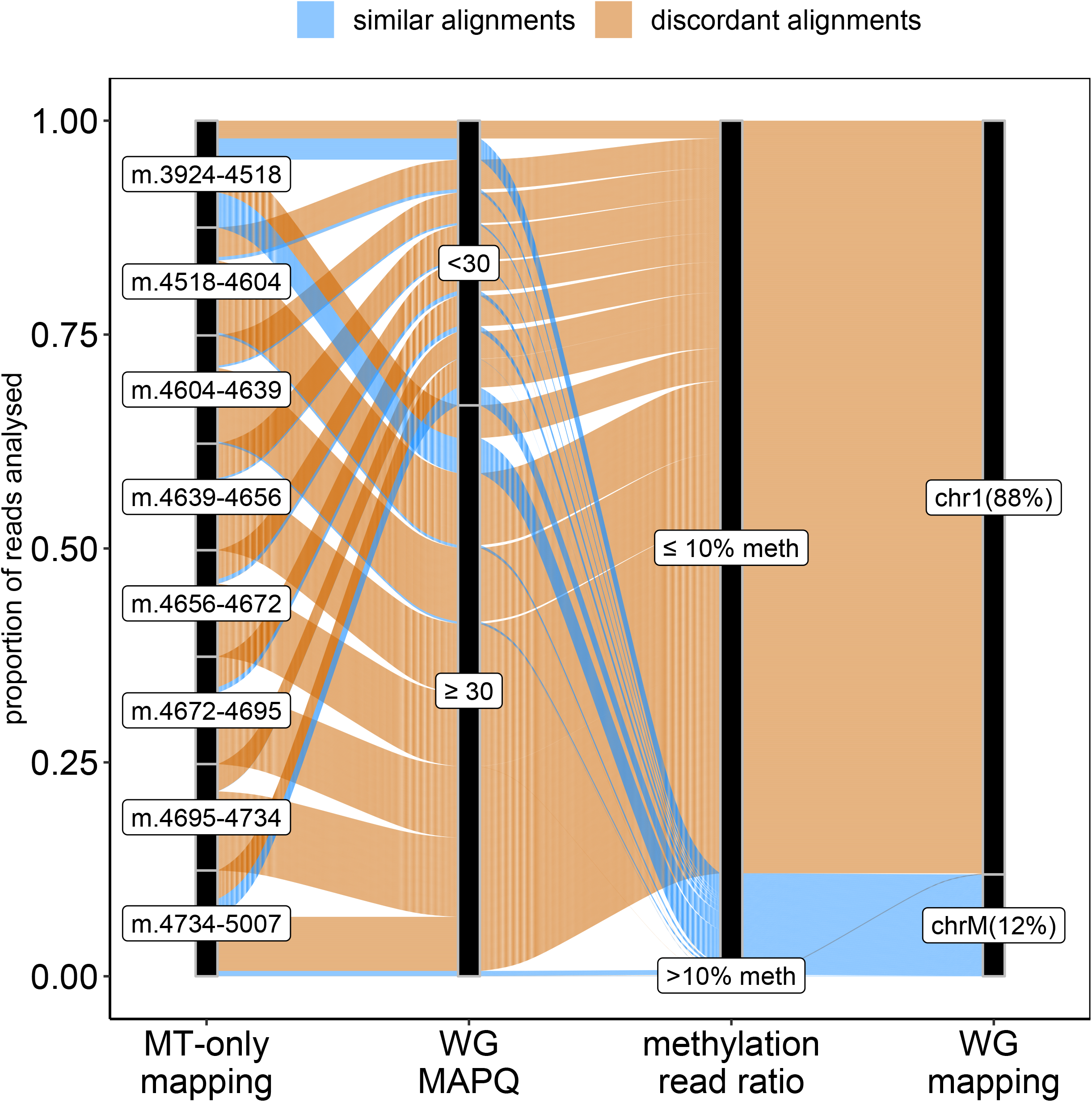
Differences in read mapping between MT-only and WG-alignment strategies. The alluvial plot traces the mapping of the reads mapped with MT-only on the positions covered by less than 1,000 reads in WG-alignment. Concordant read alignment is shown in blue and discordant alignment is shown in orange. The majority (88%) of the 8,082 reads mapped on chromosome 1 (chr1) and only 12% mapped on mtDNA (chrM) in the WG-alignment. MAPQ: mapping quality determined using Bismark (Phred score). Methylation read ratio: ratio of non-converted cytosines to converted cytosines in any given read. WG-alignment: chromosome where the reads mapped with the WG-alignment strategy.

### mtDNA harbours a single methylation hotspot in the heavy-strand promoter

Having established a framework for mtDNA analysis from WGBS data, we next assessed the mtDNA methylation landscape based on the MT-only alignment. All mtDNA cytosines were included in the analysis, with the exception of those within the first and last 125 positions of the rCRS reference, which were removed to avoid lower mapping qualities and depth caused by the linearisation of the mtDNA reference sequence. This resulted in a total of 7,218 cytosines. Bisulfite conversion efficiency, as estimated by the unmethylated Lambda phage spiked into each sample, was very high (99.64±0.01%). Thus, the contribution of non-converted unmethylated cytosines to the estimation of methylation proportions was deemed negligible. The mean methylation per sample was 0.37±0.04% (range 0.3–0.6%). The mean overall mtDNA methylation per position across our 53 samples was 0.37±0.15% (range 0.07–5.49%) and similar for both H-strand (0.38±0.19%, range 0.14–5.49%) and L-strand (0.37±0.14%, range 0.07–1.46%; Figure 3A). The three cytosine contexts showed comparable methylation scores (CpG: 0.39±0.23%, CHG: 0.36±0.14%, CHH: 0.37±0.14%). Only 21/7,218 positions showed methylation levels above 1% (Supplementary Table S5). Among these, the CpG position at m.545, situated in HSP1 showed the highest methylation levels at 5.49±0.97% (range 2.63–9.21%, Figure 3B). The cytosine at m.545 had a high depth of coverage (13,791 ±5,441) (Supplementary Table S6) and showed no correlation between coverage depth and methylation score across samples (Spearman’s: correlation ρ = 0.072, p = 0.6).

**Figure 3.**
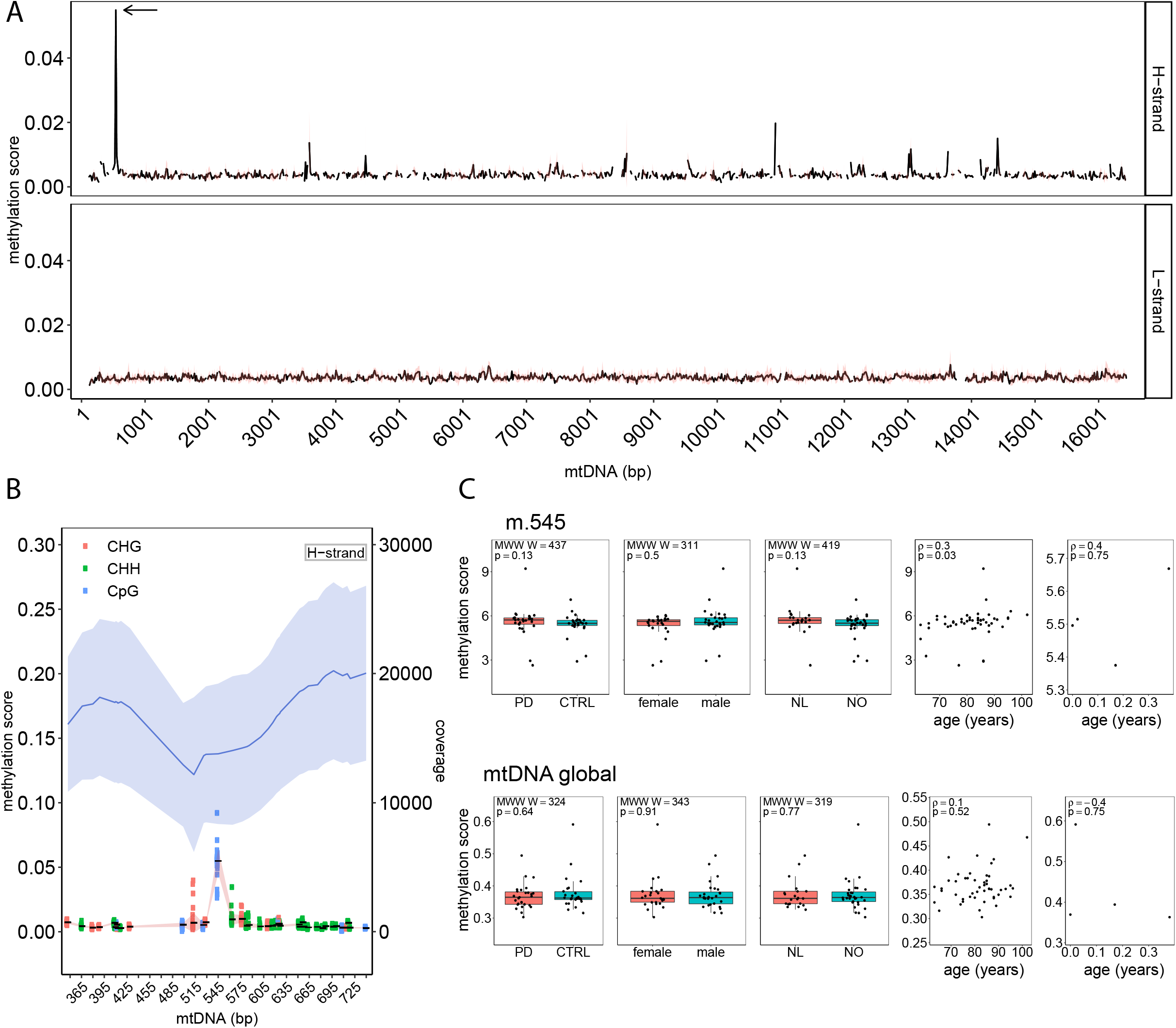
mtDNA methylation landscape in the human prefrontal cortex. A. Methylation score of mtDNA for both strands. The black line shows the average methylation score per 15-bp regions for mtDNA positions m.125-16444. The red band shows the standard deviation. Cytosine mean methylation of 15-bp regions was calculated. The arrow shows the m.545 position. B. Methylation score and depth of coverage for the mtDNA H-strand surrounding the position m.545 (m.345-745). In the upper part of the plot, mean mtDNA coverage is shown as a blue line and the light blue bands indicate the standard deviation. In the lower part of the plot, the mean methylation for each cytosine is shown as a solid black line with standard deviation indicated by the red band. The genomic context of each cytosine is indicated by different colours. C. Associations between mtDNA methylation score (global and m.545) and metadata, including disease status (PD vs controls), sex, origin of cohort (Netherlands (NL) or Norway (NO)) and age. The bar plots show no difference between the groups (Mann-Whitney-Wilcoxon test). The scatter plots show a significant correlation between m.545 methylation and age in adult samples (Spearman ρ=0.3, p=0.03) while showing no significant correlation between age and the global methylation scores.

The levels of m.545 methylation (but not global mtDNA methylation) showed a positive correlation with age in the adult group (Spearman’s: ρ = 0.3, p = 0.03, Figure 3c). This correlation survived after removal of extreme values (ρ =0.3, p = 0.01), suggesting it was not driven by outliers in the data. We found no association between mtDNA levels (global or m.545) and subject sex, or disease status (i.e., PD vs. control; Supplementary Table S1, Figure 3C).

### m.545 methylation is validated by bisulfite-independent methods

Next, we sought to validate the m.545 methylation with a sequence-specific and methylation-dependent restriction digestion of mtDNA followed by detection by qPCR or fluorescent PCR-fragment analysis. The restriction enzyme MspJI recognises a specific DNA sequence covering m.545 in a methylated state (Figure 4A). Therefore, m.545 methylation directly affects DNA restriction by MspJI at this specific site. To assess the suitability of MspJI, we digested hybridised DNA-oligonucleotide sequences mimicking the different methylation states of the MspJI restriction site, namely non-, hemi- or fully methylated. As shown in Figure 4A, the hemi- and fully methylated hybrids were efficiently digested by incubation with MspJI (including the hybrid mimicking methylation of only m.545 on the H-strand), while the non-methylated hybrid remained uncut. This DNA sequence is also digested by MspJI if only m.542 (C on the opposite L-strand) is methylated. This methylation was, however, found only at very low levels in our WGBS analyses.

**Figure 4.**
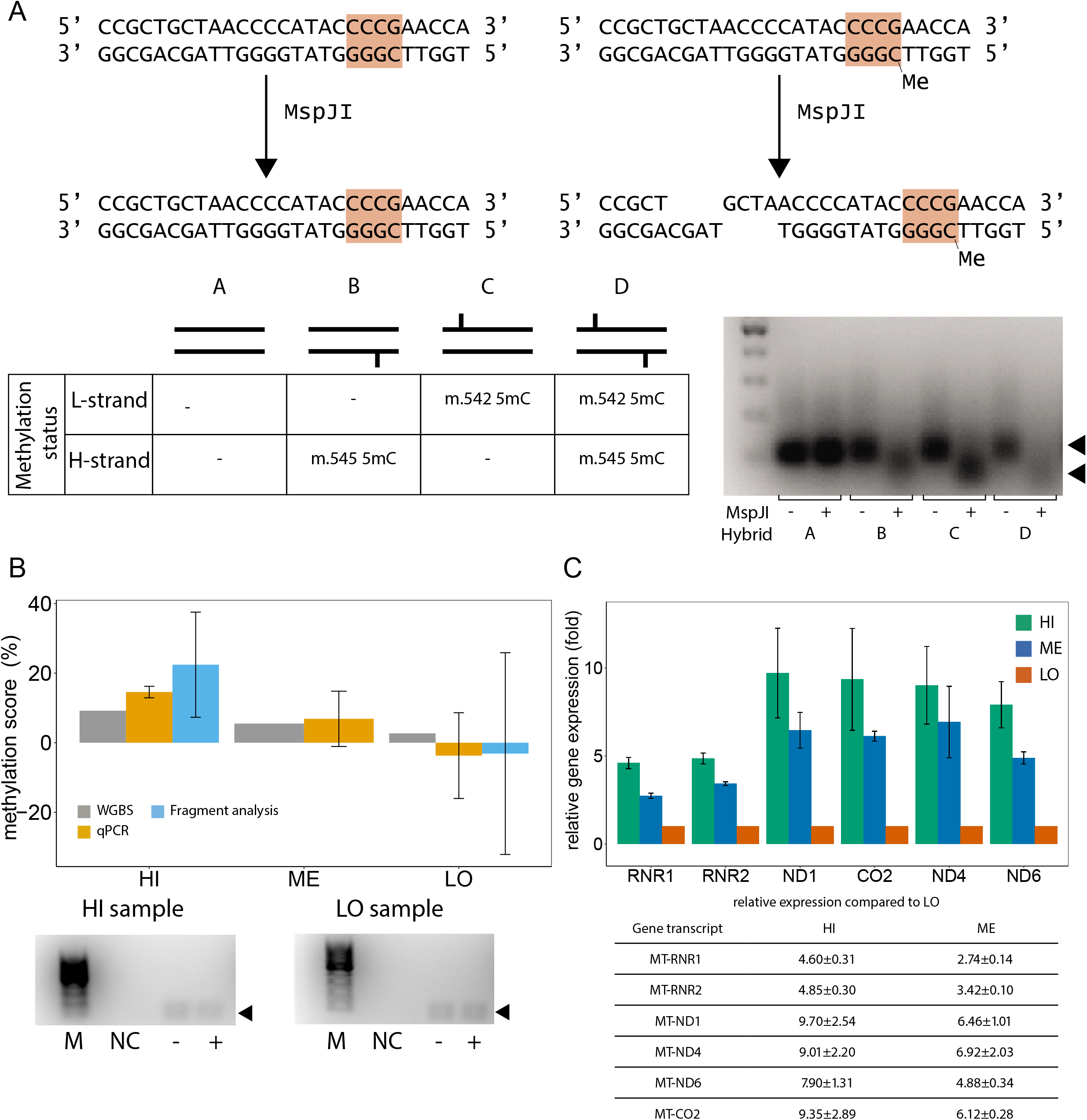
Bisulfite-independent analyses confirm m.545 methylation and indicate an effect on mitochondrial transcription. A. Upper panel: MspJI recognises mtDNA m.545 in a methylated state and only then cuts the mtDNA molecule upstream of the recognition site (orange). Lower left panel: schematic overview over synthetic oligonucleotide hybrids representing the MspJI restriction site in non-, hemi-, or fully methylated state. Lower right panel: Synthetic oligonucleotide hybrids harboring one or two methylated cytosines in the MspJI recognition site are digested by incubation with MspJI. Left lane: 100bp marker. B. Combination of MspJI restriction digestion and quantitative PCR analysis confirms mtDNA m.545 methylation Upper panel: mtDNA m.545 methylation scores obtained by WGBS (grey), qPCR (orange) and fluorescent PCR fragment analysis by capillary electrophoresis (cyan) for representative samples with high (HI), medium (ME) and low (LO) m.545 methylation scores. Lower panel: fluorescent PCR products amplified after incubation of template mtDNA with (+) or without (-) MspJI and used for quantification in are shown. NC: negative control. M: 100bp marker. C. Mitochondrial gene expression levels in samples with high, medium, and low m.545 methylation score. The means of six replicates are shown. Error bars: standard deviation.

We then isolated and enriched mtDNA from frontal cortex tissue samples with high (HI, 9.2%), medium (ME, 5.5%) and low (LO, 2.7%) WGBS methylation scores for m.545. MspJI restriction digestion and subsequent qPCR analysis comparing samples incubated with and without MspJI showed methylation levels in the range of the *in silico* data obtained by WGBS using the MT-only alignment strategy (Figure 4B, Table 1).

In a parallel approach, analysis of fluorescent-labelled PCR products of the samples with high and low methylation scores showed comparable values and further confirmed the expected range of methylation (Figure 4B, Table 1). DNA sequencing of the amplified product in both PCR approaches and a subsequent BLAST (48) search confirmed a 64bp-long DNA fragment originating from m.511-575, devoid of contamination from resembling sequences (i.e. NUMTs).

### m.545 methylation levels correlate with mitochondrial transcription

Finally, we sought to determine whether m.545 methylation is associated with mtDNA transcription. *In vitro* studies have shown that the region surrounding position m.545 (m.532-553) is a binding site for mitochondrial transcription factor A (TFAM) (49) and that increased methylation of this region increases TFAM binding and mtDNA transcription (50). To determine whether m545 methylation is associated with mtDNA transcription in the human brain, we therefore measured the expression levels of several mitochondrial genes and assessed their correlation with m.545 methylation for the same three samples as before. The reference sample was the m.545 lowest methylation sample determined by WGBS. The expression of HSP1-dependent genes (*MT-RNR1, MT-RNR2*) positively correlated with the m.545 methylation score (Spearman’s correlation ρ = 0.97, p = 0.001) (Figure 4C). Interestingly, the same trend was observed for non HSP1-dependent genes (*MT-ND1, MT-CO2, MT-ND4, MT-ND6*), both independently and collectively (ρ = 0.85 p < 10^-6^, Figure 4C). This suggests that m.545 methylation may regulate the transcription of the entire H-strand and possibly also the L-strand.

Since mitochondrial transcripts levels may be influenced by mtDNA copy number (51), we estimated mtDNA copy number for each sample using the coverage depth of the WGBS dataset. The calculated mean mtDNA copy number per cell was 2,440±523 copies. No correlation was found between mtDNA copy number and transcript fold change (Spearman’s ρ = −0.21, p = 0.1) or m.545 methylation score (Spearman’s ρ = 0.12, p = 0.4).

## DISCUSSION

We show that mtDNA cytosine methylation levels in human prefrontal cortex are generally very low, with the exception of a CpG hotspot in position m.545 in the HSP1 region. This finding is supported by multiple lines of evidence. First, the m.545 methylation was supported by a high number of reads (Figure 3B), showed no correlation with sequencing depth, and was detected at similar levels independent of the alignment strategy (Supplementary Table S3). Second, the m.545 methylation robustly replicated across 53 samples from two independent cohorts originating from different populations. Third, the m.545 methylation was validated by bisulfite-independent methodologies.

The levels of the m.545 methylation showed a mild but significant increase with age in the adult samples. Age-related changes are typically seen also in nuclear DNA methylation (52), although their physiological role remains largely unknown. However, no significant association was found between mtDNA methylation levels and disease status. Thus, brain mtDNA methylation is unlikely to be involved in PD. Our results are in contrast to those of a previous study, reporting loss of mtDNA methylation levels in the D-loop in bulk *substantia nigra* tissue of individuals with PD (19). This may be due to the different anatomical area being studied. However, since the *substantia nigra* is characterised by severe neuronal loss and gliosis in PD, it cannot be excluded that the reported mtDNA methylation changes reflect altered cell composition. Since the prefrontal cortex is not affected by severe neuronal loss in PD, our results are substantially less likely to be biased by cell-composition differences between patients and controls.

The underlying mechanisms regulating mtDNA methylation are not known. Several reports (21, 53–56) have indicated the presence of nuclear DNA methyltransferases (DNMTs) also in the mitochondria. These were, however, contradictory with regard to which DNMT isoform may be mitochondrial (53, 56). Another intriguing possibility would be mitochondrial proteins *moonlighting* as DNA methyltransferase. Such a mechanism has recently been shown for two of the three proteins that form the mitochondrial RNAse P complex responsible for tRNA processing. These two proteins form a subcomplex with RNA methyltransferase activity (57). It is possible that other such complexes exist and carry out functions that are independent of the described activities of their constituent proteins. A moonlighting activity could also explain the low level of methylation detected. Interestingly, methylation of the m.545 position to a similar low extent as our results was recently reported in human fibroblasts and cell lines (58).

Our findings suggest that m.545 methylation may influence mtDNA transcription. The transcription of HSP1-dependent genes is partly mediated by TFAM binding on regions m.425:446 and m.532-553 (49, 50). Furthermore, it has been shown that methylation of m.545 and three other cytosines (m.524, m.525, m.544) within those regions increase TFAM binding affinity *in vitro* and induce transcription of HSP1 controlled genes (50). In line with these *in vitro* observations, we show that the levels of m.545 methylation are strongly correlated with the levels of HSP1-directed mtDNA transcripts. Interestingly, however, the m.545 methylation in our samples correlated with the expression of multiple mtDNA transcripts originating from both strands. This observation suggests that the m.545 methylation may be regulating the expression of all mtDNA genes in the human brain. Indeed, it has been proposed that HSP1 may be regulating the transcription of the entire H-strand (59, 60), whereas HSP2 has been reported to be functional *in vitro* (61) but inactive *in vivo* (5).

Finally, we propose a framework for accurate mtDNA methylation assessment from WGBS data, including recommendations for the alignment strategy and minimum coverage depth. We show that alignment of mtDNA reads from WGBS data to the whole-genome reference is prone to substantial misalignment errors caused by NUMTs. Specifically, two types of misalignment errors occur. First, reads originating from the mtDNA are attracted to NUMT sequences, mainly on a region of chromosome 1 rich in mitochondrial pseudogenes. This results in a marked drop in coverage along specific mtDNA regions increasing the dispersion and hence severely compromising the resolution of methylation estimates. In addition, since bisulfite conversion of unmethylated cytosines to thymines decreases sequence similarity to the reference, it is highly likely that the loss of methylated/unmethylated mtDNA reads will be non-uniform, hence biasing the relative mtDNA methylation estimation. Second, reads originating from nuclear DNA can be erroneously aligned to the mtDNA, potentially causing overestimation of mtDNA methylation scores, since cytosine methylation is much more common in the nucleus. However, since the sequencing depth of nuclear DNA is several orders of magnitude lower than that of mtDNA in a typical WGBS experiment, misalignment of nuclear reads to mtDNA is, on its own, unlikely to represent a major source of methylation bias. These effects, combined with other technical artifacts (e.g., imperfect bisulfite conversion, low sequence complexity in the alignment stage), will be further amplified in low coverage regions. We find that the most accurate approach consists in aligning to the mtDNA reference chromosome only. This yielded both a high and uniform mtDNA coverage and a high specificity in the alignments for all samples, allowing for confident methylation assessment.

Irrespective of which alignment strategy is employed, the risk of detecting false positive methylation increases with low depth of coverage. The binomial nature of methylation proportions estimated using sequencing reads implies that the error will be asymmetrical in the extremes of methylation (i.e. 0 and 1), with an increasing number of positions in the sample deviating from the mean values as the depth approaches zero. In order to confidently estimate methylation levels when the methylation levels are near zero (or one) a high depth of coverage is a prerequisite. As a rule of thumb, to confidently assess methylation scores as low as 1%, we require 10 supporting methylated reads for the “minor allele”, and thus a total depth of ≥ 1,000X. This is in line with the notion that a confidence interval estimation of the probability of success in a binomial distribution should only be used when M = n × min (p,1-p), where p is the methylation score and M should be at least 5 or 10 (62).

One previous study in human frontal cortex (12) reported high levels of non-CpG methylation in mtDNA with a strong predilection for the L-strand. However, this study had a surprisingly low coverage of the L-strand (mean 48.5±20.2, range 5-125), which was ~57 times less than the H-strand in their experiment. While a slightly lower coverage of the L-strand is expected in bisulfite sequencing, due to its C-rich sequence predisposing to backbone breakage (63), this is usually in the order of ~2/3 of the coverage of the H-strand (64), as indeed seen in our data. The large discrepancy between the coverage of the two mtDNA strands in the study by Dou *et al*. indicates technical limitations with the analysis. Moreover, our findings show that estimating mtDNA methylation at this coverage range is highly prone to false positive results and substantial overestimation of any real methylation events. We therefor believe that these results should be considered non-informative with regard to mtDNA methylation.

Taken together, our findings provide robust evidence that mtDNA cytosine methylation is very low in the human prefrontal cortex, with the exception of a few CpG positions. Most prominent among these is the m.545 within the heavy-strand promoter, which may be involved in regulating the expression of mtDNA encoded genes. Moreover, we identify an important source of error in mtDNA methylation assessment from WGBS data and propose a robust framework to circumnavigate this bias and improve the accuracy of this type of analysis. Further work is warranted to explore the role of m.545 methylation in mitochondrial biology, including its relationship to mitochondrial transcription.

## DATA AVAILABILITY

The datasets and scripts required to reproduce the results of these analyses are available in the Neuromics Group repository, https://git.app.uib.no/neuromics/mtdna_methylation

## SUPPLEMENTARY DATA

Supplementary Data are available at NAR online.

## AUTHOR’S CONTRIBUTION

RG participated in the study conception and design, performed the computational analyses, and designed and performed the bisulfite-independent experiments. CD participated in the study conception and design and supervised the bisulfite-independent analyses. GA and OBT contributed biological material. GSN participated in the study conception and design, performed the pre-processing of the WGBS data, and supervised the bioinformatics analyses. CT conceived, designed and directed the study, contributed to data analyses and interpretation and acquired funding for the study. RG wrote the manuscript with the active input and participation of CD, GSN and CT. All authors have read and approved the manuscript.

## ETHICS DECLARATION

Ethics approval for these studies was obtained from our regional ethics committee (REK 2017/2082, 2010/1700, 131.04). Consent for publication was provided by all participants. The authors declare that they have no competing interests.

