## Supplementary material for "Ultra-deep whole genome bisulfite sequencing reveals a single methylation hotspot in human brain mitochondrial DNA": suppltable1.docx

**Supplementary Table 1. Comparison of the main outcomes for the three alignment strategies per strand.**

| **Alignment strategy, depth filter, (strand)** | **mean depth ± SD (range)** | **mean methylation  ± SD (range)** | **mean methylation ± SD (CpG / CHG / CHH)** | **number of cytosines methylated > 1% (mean methylation  ± SD)** | **context of cytosines methylated > 1% (CpG / CHG / CHH)** | **m.545 mean methylation  ± SD (range)** |
| --- | --- | --- | --- | --- | --- | --- |
| **MT-only,  No depth filter,  (L)** | 12451±3469 (4535 - 24024) | 0.37±0.14% (0.07 - 3.3) | 0.38±0.1 / 0.35±0.11 / 0.37±0.15 | 8 (1.46±0.77) | 0 / 1 / 7 | N.D. |
| **MT-only,  No depth filter,  (H)** | 27055±4832 (12179 - 37696) | 0.38±0.19% (0.14 - 5.49) | 0.4±0.3 / 0.38±0.18 / 0.38±0.14 | 17 (1.65±1.06) | 6 / 7 / 4 | 5.49±0.97%  (2.63 - 9.21) |
| **MT-only,  No depth filter,  (both strands)** | 16719±7711 (4535 - 37696) | 0.37±0.16% (0.07 - 5.49) | 0.39±0.23 / 0.36±0.15 / 0.37±0.15 | 25 (1.59±0.97) | 6 / 8 / 11 | 5.49±0.97%  (2.63 - 9.21) |
| **WG,  No depth filter,  (L)** | 10999±3992 (81 - 23082) | 0.36±0.16% (0.07 - 3.33) | 0.34±0.12 / 0.34±0.13 / 0.37±0.16 | 12 (1.39±0.63) | 0 / 2 / 10 | N.D. |
| **WG,  No depth filter,  (H)** | 25344±5671 (9175 - 36105) | 0.37±0.19% (0.1 - 5.49) | 0.38±0.31 / 0.38±0.18 / 0.37±0.14 | 18 (1.6±1.04) | 6 / 7 / 5 | 5.49±0.98% (2.52 - 9.21) |
| **WG,  No depth filter,  (both strands)** | 15192±7952 (81 - 36105) | 0.37±0.17% (0.07 - 5.49) | 0.36±0.23 / 0.36±0.16 / 0.37±0.16 | 30 (1.51±0.89) | 6 / 9 / 15 | 5.49±0.98%  (2.52 - 9.21) |
| **two-stage,  No depth filter,  (L)** | 5093±4943 (5 - 20423) | 0.73±1.06% (0.04 - 33.48) | 0.75±1.14 / 0.69±0.78 / 0.73±1.09 | 859 (2.21±1.95) | 79 / 79 / 701 | N.D. |
| **two-stage,  No depth filter,  (H)** | 15894±10664 (423 - 33981) | 0.53±0.46% (0.05 - 5.47) | 0.54±0.53 / 0.53±0.44 / 0.52±0.43 | 220 (1.59±0.65) | 52 / 47 / 121 | 5.47±0.95%  (2.54 - 8.86) |
| **two-stage,  No depth filter,  (both strands)** | 8276±8659 (5 - 33981) | 0.67±0.93% (0.04 - 33.48) | 0.65±0.89 / 0.61±0.65 / 0.68±0.98 | 1079 (2.08±1.79) | 131 / 126 / 822 | 5.47±0.95% (2.54 - 8.86) |
| **MT-only,  Depth filter = 1000,  (L)** | 12454±3466 (4611 - 24024) | 0.37±0.14% (0.07 - 1.46) | 0.38±0.1 / 0.35±0.11 / 0.37±0.14 | 5 (1.17±0.22) | 0 / 0 / 5 | N.D. |
| **MT-only,  Depth filter = 1000, (H)** | 27056±4832 (12179 - 37696) | 0.38±0.19% (0.14 - 5.49) | 0.4±0.3 / 0.38±0.18 / 0.38±0.13 | 16 (1.65±1.09) | 6 / 7 / 3 | 5.49±0.97%  (2.63 - 9.21) |
| **MT-only,  Depth filter = 1000,  (both strands)** | 16723±7710 (4611 - 37696) | 0.37±0.15% (0.07 - 5.49) | 0.39±0.23 / 0.36±0.14 / 0.37±0.14 | 21 (1.54±0.98) | 6 / 7 / 8 | 5.49±0.97%  (2.63 - 9.21) |

Mean depth is defined for all positions in each dataset. The depth filter
(when applied) excluded a position if not supported by at least 1000 reads. Cytosine contexts are defined such as CpG: a C followed by a G; CHG and CHH refer to a triplet where H stands for a nucleotide different from G
(i.e. A, C, T). N.D.: non defined. SD: standard deviation.
