## Supplementary material for "Ultra-deep whole genome bisulfite sequencing reveals a single methylation hotspot in human brain mitochondrial DNA": table1.docx

| **Sample** | **HI** | **ME** | **LO** |
| --- | --- | --- | --- |
| **WGBS** | 9.2% | 5.50% | 2.7% |
| **qPCR** | 14.6±1.65% | 6.86±7.94% | -3.7±12.32% |
| **Fragment analysis** | 22.42±15.1% | NA | -3.12±29% |

**Table 1. m.545 methylation scores obtained from WGBS and bisulfite independent methods.** NA : not available, no medium-methylated sample has been included in the fragment analysis experiment.
