## Supplementary figures and images for "Ultra-deep whole genome bisulfite sequencing reveals a single methylation hotspot in human brain mitochondrial DNA"

### supplfig1.pdf

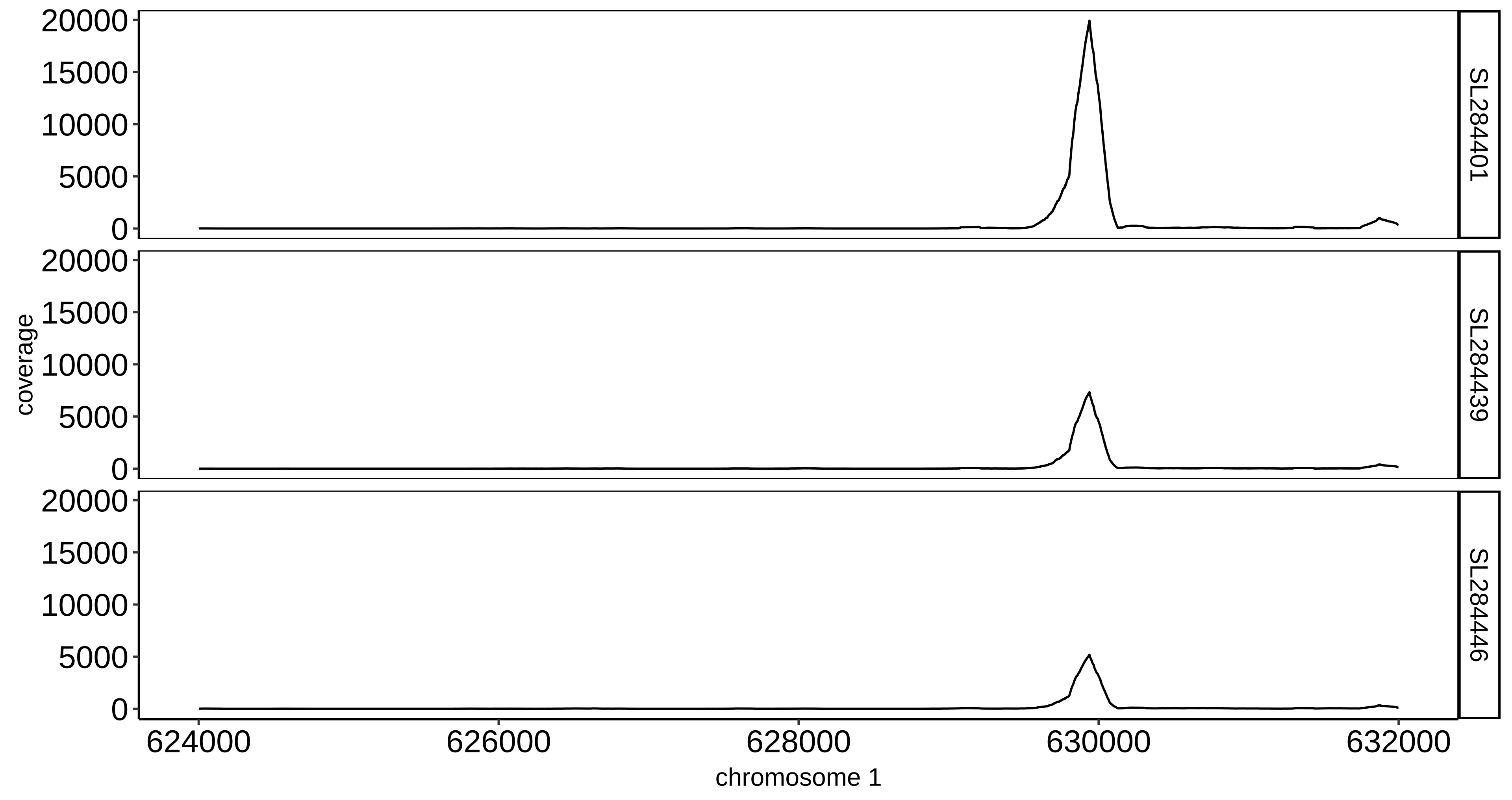

### supplfig2.pdf

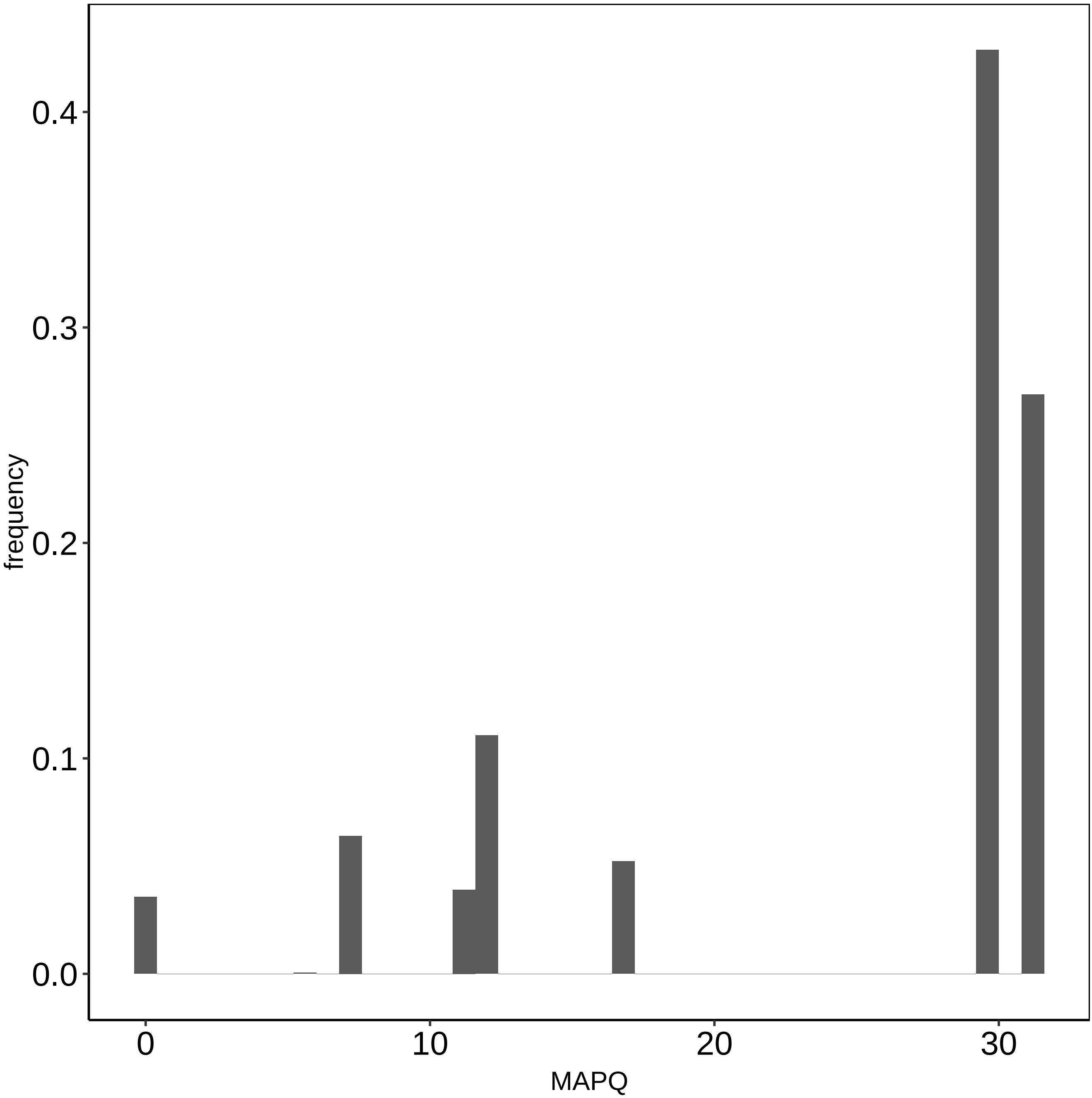

### supplfig3.pdf

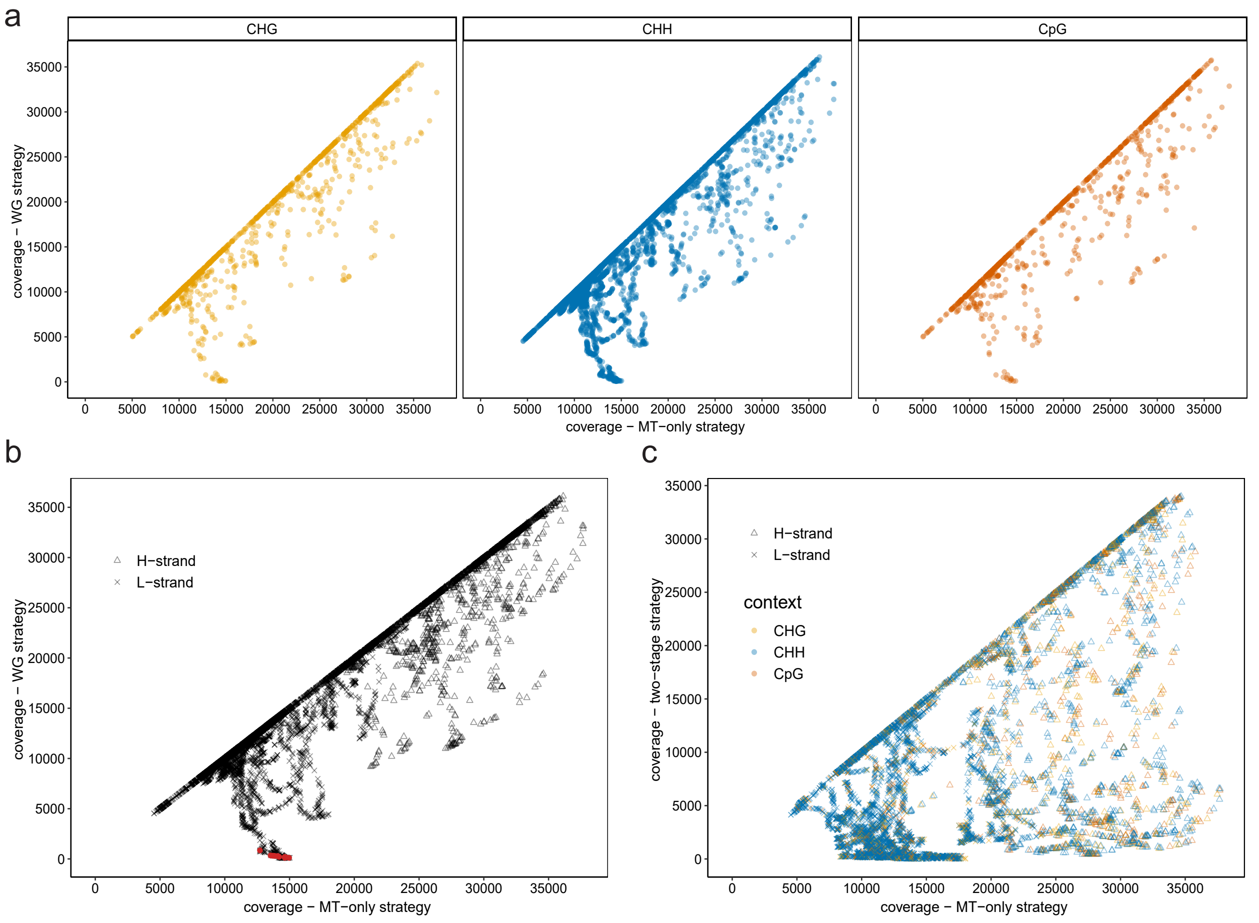
